# Glymphatic system health in early Alzheimer’s disease and its relationship to sleep, cognition and CSF biomarkers

**DOI:** 10.1101/2024.10.14.618324

**Authors:** Nicola Jane Ray, Natalia Cullell, Oliver Clark, Ariane Delgado-Sanchez, Giovanni Caruana, Andrea Elias-Mas, Julia Ferrer, Mariateresa Buongiorno, Judith Castejon, Gonzalo Sánchez-Benavides, Esther Granell, Sonia Correa-Muller, Jerzy Krupinski

## Abstract

**BACKGROUND:** The glymphatic system is thought to facilitate waste clearance from the brain during sleep. Impairment in this system may underpin the elevated deposition of pathological proteins in neurodegenerative conditions like Alzheimer’s disease (AD). Putative glymphatic system activity has been measured with contrast-enhanced, serial MRI, revealing slower clearance in people who are sleep deprived. It is important that these methods are used to understand changes to glymphatic function in people with early-stage AD.

**METHODS:** Twenty-four individuals with mild cognitive impairment were recruited. N=20 had CSF biomarker data, with 16 meeting criteria for AD positivity (AD+). Participants underwent polysomnography, cognitive testing and serial T1 MRI with intravenous gadolinium-based contrast agent, diffusion tensor imaging along the perivascular space (DTI-ALPS index), and core AD CSF biomarkers collection. Rate (over 24hrs) and efficiency (the amount of tracer cleared after 28hrs relative to uptake after 4hrs) of GCBA clearance were measured.

**RESULTS:** Faster/more efficient GBCA clearance was associated with shorter sleep latency. In AD+ participants faster 24hr clearance of GBCA was associated with a lower ratio of Aβ1–42/Aβ1–40 in the CSF. In addition, better clearance efficiency was associated with greater levels of Aβ1-40, lower levels of Aβ-1-42, and a smaller ratio of Aβ1–42/Aβ1–40 in the CSF. Higher DTI-ALPS indicated better cognitive performance and, unexpectedly, higher tau levels. However, it was not associated with GCBA clearance.

**CONCLUSIONS:** We show, for the first time in humans, that glymphatic system function is associated with AD-related changes to sleep, cognition and core AD biomarker concentrations in CSF. However, for AD biomaker concentrations, these relationships are not in the expected direction: higher concentrations and lower Aβ1–42/Aβ1–40 ratio were associated with faster or more efficient clearance of GCBA. We suggest that increased neurodegeneration in those with elevated levels of tau and Aβ1–40 may paradoxically increase glymphatic activity locally to brain regions with more atrophy relative to less, but not in a way that improves sleep or cognition. In fact, larger extracellular spaces may exacerbate the spread of tau via the glymphatic system and therefore accelerate the progression of AD.

## Background

The discovery of the glymphatic system^1^ and functional meningeal lymphatic vessels (enabling molecular egress from CSF to extracranial lymph)^2–4^ has been closely followed by hypotheses that implicate impairments in these clearance routes within pathophysiological pathways underpinning neurodegenerative conditions, including Alzheimer’s disease (AD) ^4,5^.

Research in animals using fluorescent tracers reveals that the glymphatic system supports transport of cerebrospinal fluid (CSF) into the brain parenchyma within subarachnoid spaces, where exchange between CSF and interstitial fluid (ISF) occurs. In humans, we can observe the glymphatic system via contrast-enhanced magnetic resonance imaging (MRI). Indeed, serial MRI following intrathecal gadobutrol injection has been used to visualise CSF-ISF exchange in humans, which has confirmed that the glymphatic system is active during sleep, and that overnight sleep deprivation reduces glymphatic activity ^1,6^.

Non-invasive methods to index glymphatic system function are also proposed. Notably, diffusion tensor imaging (DTI), using diffusion-weighted images (DWI), provides a metric that indirectly analyses water diffusivity along perivascular spaces (PVS) (the DTI-ALPS index^7^). There has been a sharp increase in the number of studies using this index despite no direct evidence that it is a marker of glymphatic system function. Indeed, it is questioned whether movement of water in white matter can feasibly represent mechanisms that underpin the clearance of large-molecular solutes from the cortex ^8^. Yet, relationships are increasingly reported between the ALPS-index and sleep ^9^, cognitive status ^10^, CSF biomarker concentrations ^10,11^, and intrathecal gadolinium-based contrast agent (GBCA) clearance ^12^.

To further examine the role of glymphatic system function in the development of AD, it is necessary to examine its relationship with CSF biomarker concentrations, sleep and cognitive performance in people at the earliest disease stages. In previous work, we found people with AD pathology, sleep disturbances, and mild-to-moderate cognitive impairment developed severe side effects from intrathecal administration of gadobutrol, not previously observed in healthy samples, which necessitated a discontinuation of the study ^13^. The participants who finished the protocol displayed persistent tracer enrichment in the cortex and white matter, suggesting that brains with sleep impairment and AD pathology have poor glymphatic function, and cannot clear the contrast tracer efficiently.

Recent work has instead used intravenously administered gadolinium-based contrast agent (GBCA) to study the human glymphatic system ^14–18^. However, the route through which GBCA may reach the CSF has not been established. It is speculated that intravenously administered GBCA enters periarterial spaces in people with disrupted blood–brain barrier (BBB). The contrast would then flow into the interstitial compartment, to be subsequently cleared via meningeal lymphatics ^18^. In those without loss of BBB integrity, intravenous GBCA has been suggested to enter CSF from the choroid plexus, cranial nerve sheath, or dural vessels ^18,19^. Recently, ^20^ used real-IR MRI to observe rapid spreading enhancement around bridging veins located within the subarachnoid space in healthy participants, but without evidence for entry of GBCA through the choroid plexus. While tracer-related enhancement was predominantly observed in older individuals, the authors’ data implied that BBB leakage was not the cause of this. Others have suggested instead that age-related impairment in meningeal lymphatic drainage may lead to an overflow of fluid from the dura to the subarachnoid space ^21,22^.

To our knowledge, no previous research has measured glymphatic system activity via clearance of intravenously administered GBCA in people with early-stage AD. The research above leads us to expect that people with AD will have poor glymphatic function, and therefore less effective clearance of GBCA. In addition, we expect reduced GBCA clearance to be associated with worse sleep and cognitive symptoms. However, there is likely to be a more complex relationship between GBCA clearance and CSF biomarker concentrations ^23^. In addition, the animal literature implies that these relationships may be particularly complex in the context of AD-related neurodegeneration^24^. Here, we aimed to capture the between-individual variation in GBCA clearance as an index of glymphatic system activity to explore its association with neuropsychological outcomes, objective sleep assessment, CSF core AD biomarker concentrations, and the DTI-ALPS index.

## Methods

### Participants

Twenty-four participants between age 65 and 75 with memory complaints and at least 0.5 in the memory domain in the Clinical Dementia Rating (CDR) who had been referred to the Cognition and Behaviour Unit at the Department of Neurology, Hospital Universitari MútuaTerrassa, (HUMT), Barcelona, Spain were recruited to the study.

Twenty-four participants had DTI and susceptibility-weighted images (SWI) images collected (N=20 of these also had CSF biomarkers, with 16 meeting criteria for AD positivity, AD+). Of the full sample, 20 participants completed serial MRI before and after intravenous GBCA injection, with 16 of these passing quality control checks (see MRI Preprocessing and Analysis). Of these, 3 did not have CSF biomarker data available (See Figure 1 for a flow chart of participants who completed each aspect of the study).

**Figure 1.**
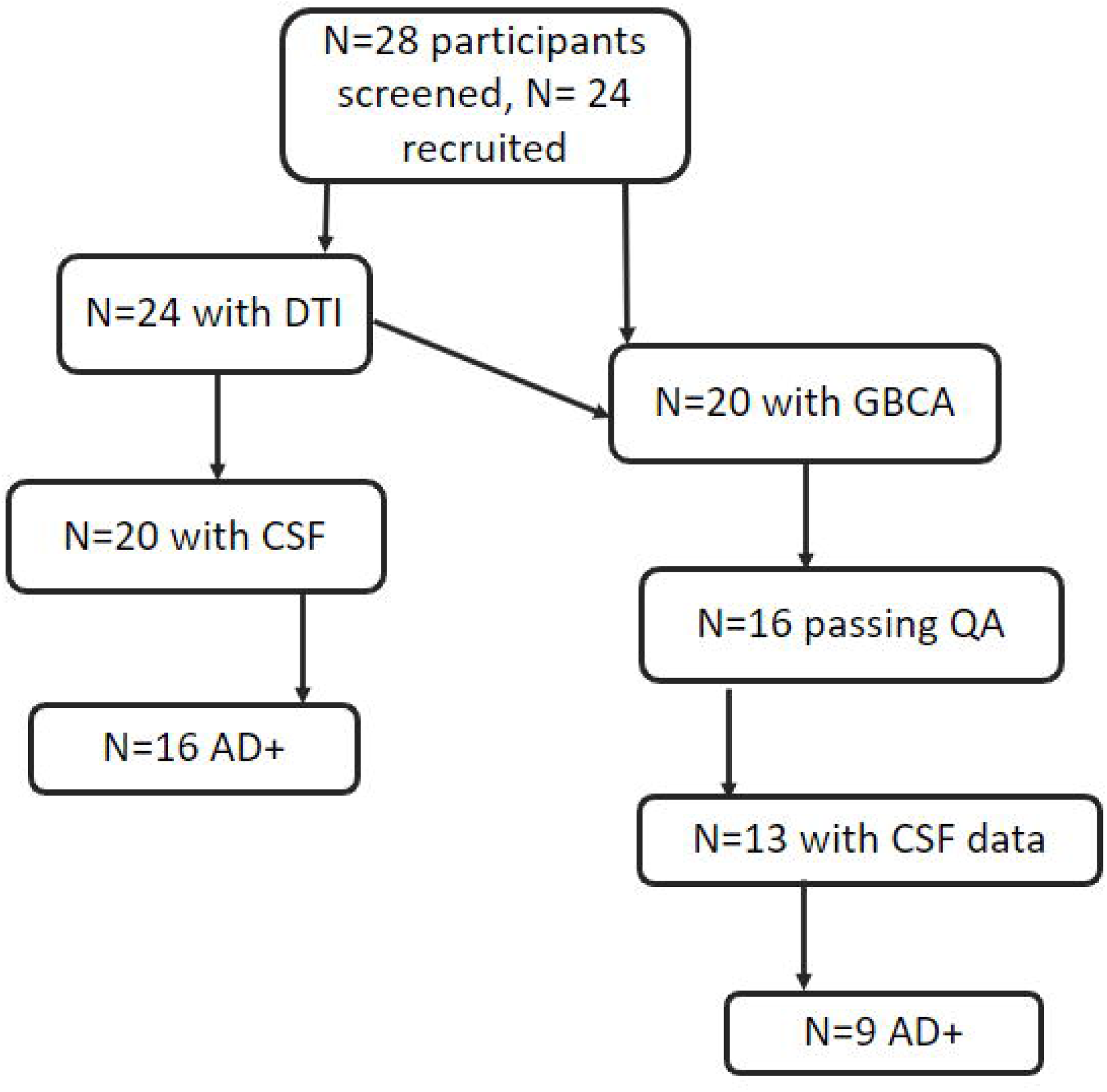
Figure 1, Flow chart of participants taking part in each element of the study. DTI, diffusion tensor imaging; CSF, cerebrospinal fluid; AD+, AD positivity via CSF biomarker levels, GBCA, gadolinium-based contrast agent; QA, quality assurance.

### Polysomnography (PSG) and sleep architecture

Participants underwent an overnight PSG at the Adsalutem sleep unit, which included videorecording, electroencephalogram (EEG), electromyography (EMG) electrodes, nasal cannula, thermistor, and pulse-oximeter. Sleep stages and respiratory events were scored according to the American Academy of Sleep Medicine standard criteria.

### Neuropsychology

Global cognitive performance was assessed using the mini-mental state exam (MMSE), and memory was assessed with the Repeatable Battery for the Assessment of Neuropsychological Status (RBANS) delayed memory index.

### Biomarkers

CSF samples, obtained by lumbar puncture, were collected in polypropylene tubes, centrifuged at 2000 rpm for 20 min, aliquoted in propylene tubes and stored at −80 °C until biochemical analysis using standard protocols. The CSF biomarkers analyses were performed by Catlab (https://www.catlab.cat/en/) and levels of Aβ1-42, Aβ1-40, total (t)-tau, and phosphorylated tau at threonine 181 (p-tau) were measured using the following: Lumipulse G β-amyloid 1-40, Lumipulse G β-amyloid 1-42, Lumipulse G pTau 181 and Lumipulse G Total Tau chemiluminescent immunoassay Lumipulse® G essay kits from Fujirebio (Fujirebio Inc. Europe, Gent, Belgium). The sensitivity and detection values were used as described by the producer and local pathological cut off values were previously described in Alvarez et al. 2018. CSF levels of Aβ1–42, Aβ1–40, total (t)-tau, and phosphorylated tau at threonine 181 (p-tau) were measured using the Lumipulse essay kits from Fujirebio (Fujirebio Inc. Europe, Gent, Belgium). We determined positivity of AD core biomarkers using local cut off values. Cut-off points to define positivity for AD were Aβ1-42/Aβ1-40 < 0.068 plus at least two others from Aβ1-42 <638 pg/mL; total (t)-tau > 404 pg/mL; p-tau 181 >52.1 pg/mL; t-tau/ Aβ42 > 0.784. In two participants who did not have lumbar puncture, AD+ was confirmed via amyloid PET scan.

### MRI acquisition

Anatomical scans were acquired using a 3T MR scanner at Radiology Department (UDIAT-Parc Taulí Sabadell) (Phillips Ingenia Elition). A standardized MR protocol was used for the acquisition, comprising of high-resolution 3D T1-weighted magnetization-prepared rapid gradient echo (MPRAGE) imaging sequence for enhanced tissue contrast, with the following settings: TR = 6.8, TE = 3.1; TI = 606.4 msec; flip angle = 8. Voxel dimensions were 1x1x1mm and the acquisition time for each scan was 4:13 min.

DWI were acquired on the same scanner with the following protocol: TR = 2.53, TE = 0.07, slice thickness = 2.2 mm, voxel size= 1.69 x 1.69 x 2.2. DWI was performed in 128 directions (diffusion b=1000 s/mm2) and in one acquisition without diffusion weighting (B0).

### MRI Preprocessing and analysis

Three of the 20 participants had artifacts on one or more of their T1-weighted scans, and were therefore removed from further analysis. One participant did not show enhancement of the T1 signal after GBCA and was removed from further analysis. We used FreeSurfer software (version 7) (http://surfer.nmr.mgh.harvard.edu/) for the segmentation, parcellation, and registration of the three T1-weighted images (using the longitudinal recon-all pipeline). Using a hybrid watershed/surface deformation procedure ^25^, non-brain tissue is removed, followed by the segmentation of the subcortical white matter and deep gray matter structures (including the hippocampus, amygdala, caudate, putamen, pallidum, and ventricles) as well as cortex ^26^. A median template was created for each participant, and the three MRI images were registered to this template via rigid transformation ^27^. The registrations were then checked manually. The segmentations and parcellations produced by recon-all for each scan were then transformed back into native T1 space to extract T1 signal intensity within each region from the native scan. Next, we used T1 signal intensity within a reference region in the posterior part of the orbit to normalize each scan according to changes in the grey-scale due to automatic image scaling, which will be different for each time point.

Glymphatic system function was indexed in two ways: the rate of clearance over 24hrs following the initial tracer uptake (4hr – 28hr/4hr); the degree of tracer cleared relative to uptake (1-(28hr-baseline)/4hr-baseline)). For the latter, a fully quantitative T-signal would give a value of 0 for no clearance of tracer, and 1 for complete clearance of the tracer. However, the normalization procedure used here (see above) does not fully normalize the T1 signal across the 3 scans, rather it minimizes the influence of scaling. To force the values from this latter calculation into a scale between 0 and 1 therefore, we have divided the outcome by 3 (since all values were below 3). This preserves the pattern of clearance across individuals, but should not be interpreted as an index that represents % of tracer cleared. Henceforth, we will describe this calculation as representing ‘clearance efficiency’.

The DWI images were processed with FSL ^28^. Images were skull stripped ^29^ and eddy current-induced distortions and subject movements were corrected ^30^. To calculate the ALPS-index FSL’s DTIFIT was used to create diffusivity maps in subject space in the x, y and z directions, as well as a colour coded vector image showing the principal diffusion tensor direction (V1). To identify the location of the PVS, SWI were spatially co-registered with the subject-space B0 image using ANTs, and superimposed on the V1 image to identify the medullary veins located perpendicularly to the PVSs and within the projection and association fibres. 4 mm spheres were placed at these locations, and diffusion in x, y and z directions spheres were extracted and used to compute the ALPS index (^17^ according to the following equation:

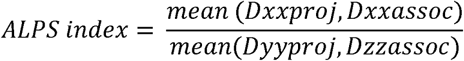

### Procedure

After recruitment at the Cognition and Behaviour Unit, and signature of the informed consent, participants underwent neuropsychological assessment. CSF collection was carried out via lumbar puncture always first thing in the morning and within one week of the polysomnography and imaging. Participants were admitted to the Adsalutem sleep unit to undergo a sleep assessment with polysomnography on their first night. Immediately following this, participants completed baseline T1-weighted and DWI scans in the morning. After 4hrs, the second T1-weighted image was collected. Participants returned to the hospital for a final scan 24hrs later. They slept freely within this time.

### Statistical analysis

This is an exploratory study to understand whether clearance of intravenously administered GBCA from the brain is associated with the early symptoms and markers of AD. Due to the presence of statistical outliers in the clearance data as determined by Grubbs test in the outliers R package^31^, we opted for a non-parametric approach. Using the *correlation* package in R we computed Bayesian and Frequentist two-tailed bi-variate rank order Pearson’s correlations between symptom markers, 24hr clearance rate and clearance efficiency (described above in ‘MRI Preprocessing and analysis’) from each grey matter brain region segmented by FreeSurfer (see above), as well as whole brain white matter and the ventricles. Due to the exploratory nature and small sample of the study, we selected an uninformed a “ultrawide” Jeffrey prior (*beta*(1,1) centred on 0) provided by the *BayesFactor* package ^32^ in R because it minimised type-1 and type-2 error-rates in simulations. Bayes Factors are a continuous index of evidence in favour of one hypothesis over another, although cut-offs are often used. In line with Jeffrey’s original suggestions, we chose a Bayes Factor of 3 as a minimal level of evidence ^33^. Effects in which the conditions p < 0.05 and BF ≥ 3 were both met were treated as evidence in favour of a non-null population effect. The correlation coefficient and Bayes Factor were then projected onto the *Desikan Killiany* and *aseg* atlases using the *ggseg* package in R. We present these outcomes uncorrected for multiple comparisons, under the proviso that all outcomes are preliminary and would need to be confirmed in hypothesis-led analyses.

Correlations between clearance, DTI-ALPS, sleep, cognitive tests were performed in the whole group (N=24 for DTI-ALPS, N=16 for GBCA clearance). Analysis of biomarker outcomes was carried out only in AD+ participants with CSF data (N=16 for ALPS, N=9 for GBCA clearance) to evaluate the glymphatic system and its relationship with CSF biomarkers in the context of AD.

## Results

Table 1 shows the demographic, clinical and sleep data for the full sample. It was notable that all participants presented with obstructive sleep apnoea, despite none being aware they were living with the condition.

**Table 1:** GBCA, gadolinium-based contrast agent; CSF, cerebrospinal fluid; TSTSpO2>60, Total sleep time with oxygenation over 60; RDI(/hour), respiratory disturbance index; WASO, Wakefulness after sleep onset; REM, rapid eye movement; MMSE, mini-mental state examination; RBANS, Repeatable Battery for the Assessment of Neuropsychological Status; DTI-ALPS, diffusion tensor images along the perivascular space. Cut-off points to define positivity for AD were Aβ1-42/Aβ1-40 < 0.068 plus at least two other from Aβ1-42 <638 pg/mL; total (t)-tau > 404 pg/mL; p-tau 181 >52.1 pg/mL; t-tau/ Aβ42 > 0.784. In two participants who did not have lumbar puncture, AD+ was confirmed via amyloid PET scan.

In Figure 2, we show 4hr and 28hr clearance of intravenously injected gadolinium from the cortical grey matter (1a), subcortical white matter (1b) and ventricles (1c), with AD-negative participants indicated by dashed lines. Figure 3 shows the 24hr clearance rate and the degree of tracer cleared from all ROIs in the participants with and without evidence of AD-related biomarkers in CSF (the outcomes across all ROIs for T1 signal enhancement after 4 and 28hrs are showing in Supplemental Figure 1). We cannot perform statistical comparisons between these groups given the low numbers, but the figures qualitatively suggest a consistent pattern in which tracer is cleared more slowly (Supplemental Figure 1a,b and Figure 3a) and less effectively (Figure 3b) in those with evidence of AD-related pathology in all brain regions.

**Figure 2.**
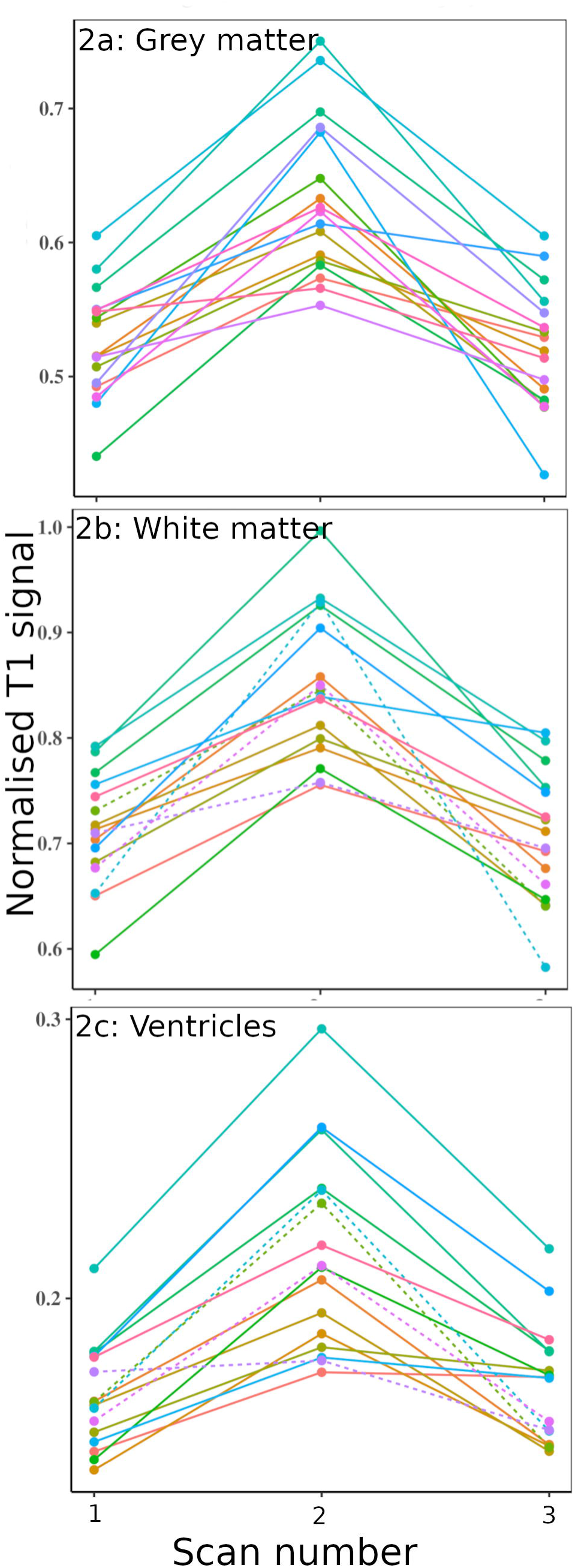
Figure 2: Normalised T1 unit at baseline and 4hr and 28h after intravenous injection of gadolinium. The averaged T1 unit from cortical grey matter is shown in 1a, subcortical white matter in 1b, and ventricles in 1c. AD-negative participants are indicated by dashed lines.

**Figure 3.**
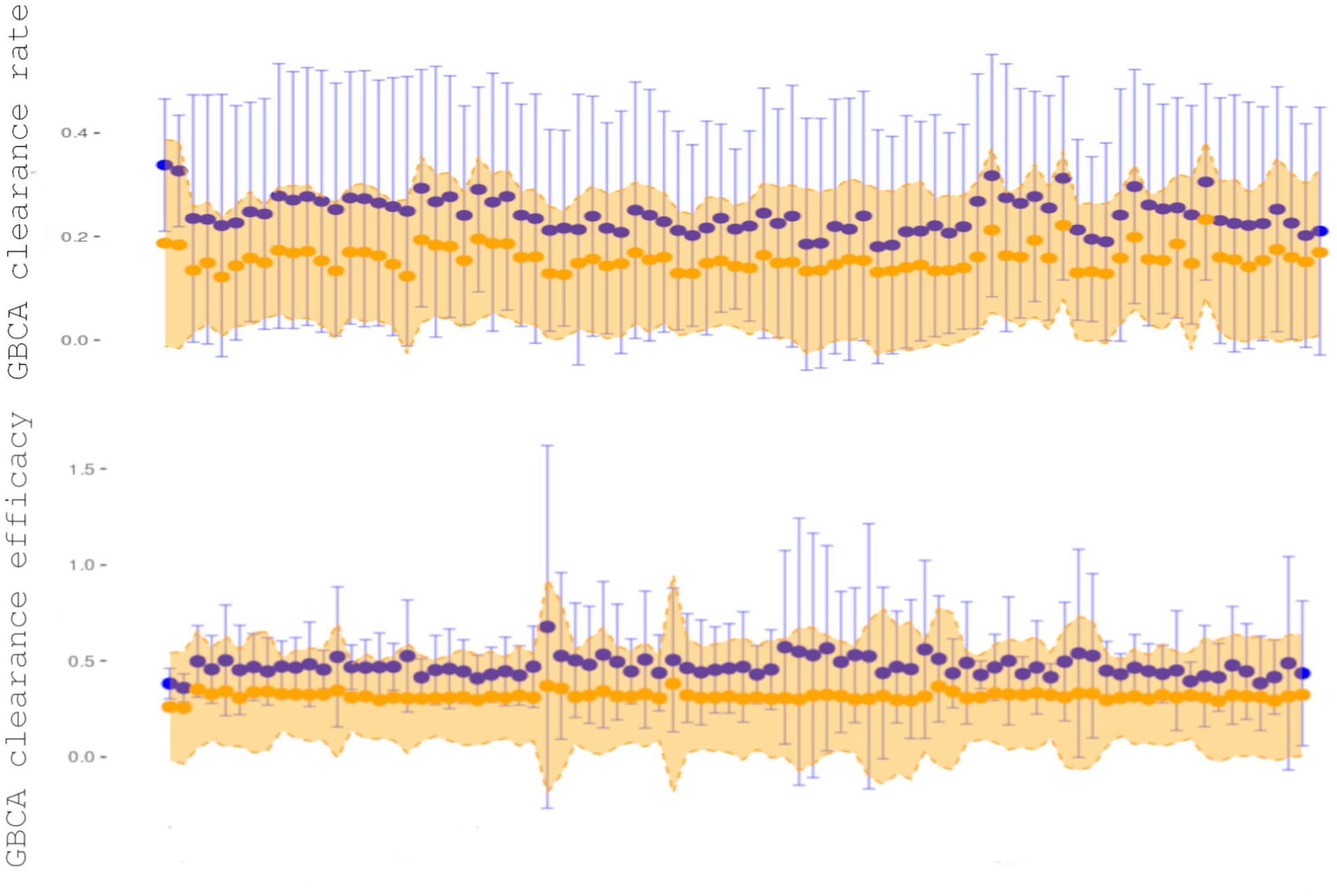
Figure 3 shows the 24hr clearance rate and the degree of tracer cleared from all ROIs in the participants with (orange) and without (blue) evidence of AD-related biomarkers in CSF. Top: the rate of clearance over 24hrs following the initial tracer uptake (28hr – 4hr/4hr), described at 24hr clearance rate. Bottom: the degree of tracer cleared relative to uptake (1-(28hr-baseline)/4hr-baseline)). For the latter, a fully quantitative T-signal would give a value of 0 for no clearance of tracer, and 1 for complete clearance of the tracer.

### Correlations between the 24hr rate of clearance of GBCA and the DTI-ALPS index, cognitive outcomes, sleep and CSF biomarker concentrations

There were no significant correlations between the DTI-ALPS index or cognitive scores and GBCA clearance rate. However, shorter sleep latency was associated with faster 24hr clearance rate in almost all brain regions analysed (Figure 4a), including white matter (cortical: Frequentist rho = -.647, p= 0.007; Bayesian: r = -.59, BF = 8.96; subcortical: Frequentist: rho = -.612, p = 0.012; Bayesian r = -.54, BF = 5.69). In the sub-sample with evidence of AD pathology, faster 24hr clearance rates in frontal and temporal cortical regions and the right amygdala were associated with a lower Aβ1–42/Aβ1–40 ratio (Figure 4b).

**Figure 4:**
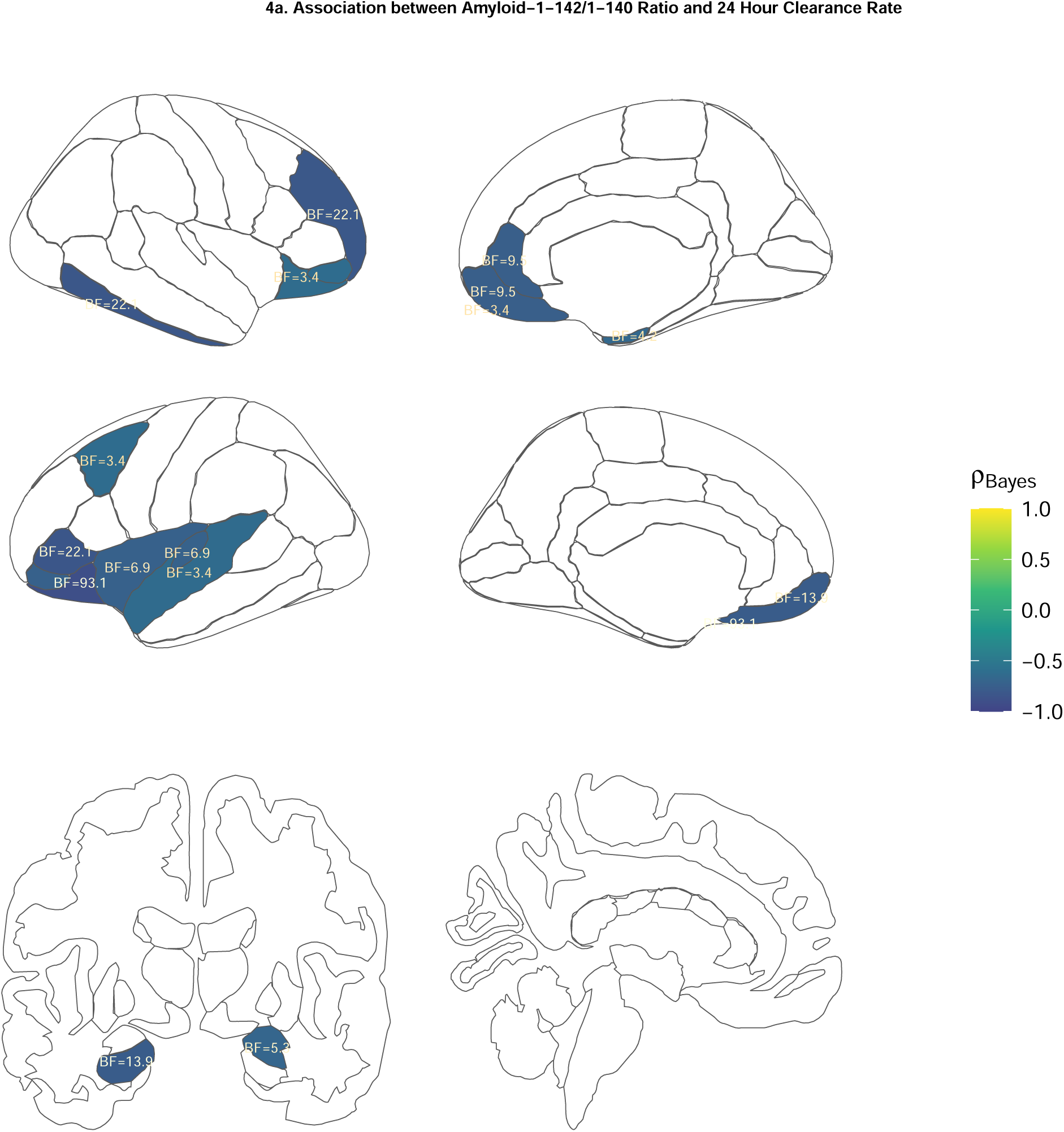
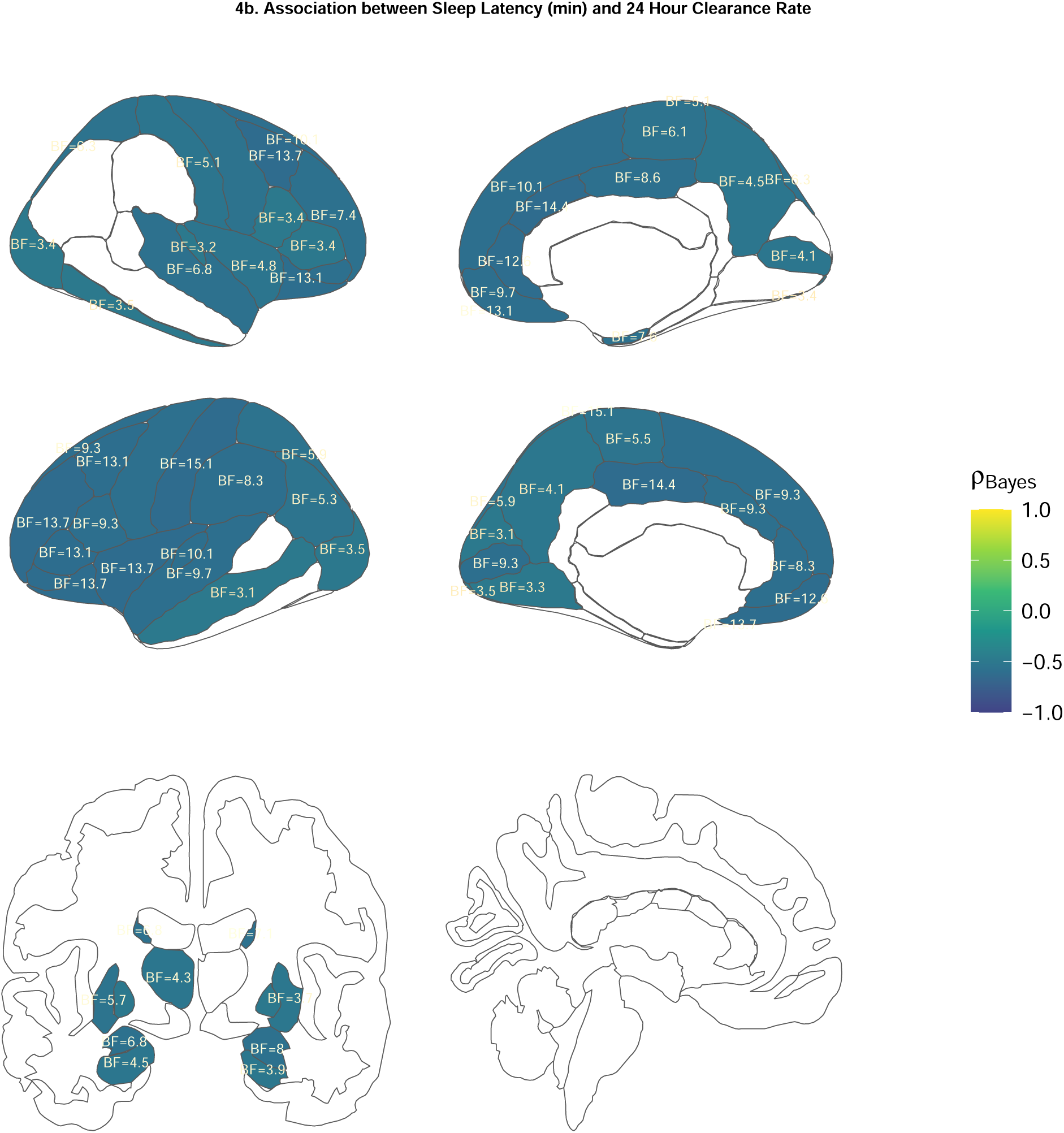
24hr clearance rate. Figure 4a, regions showing significant negative correlation between shorter sleep latency and 24hr clearance rate in all participants. Figure 4b, regions showing significant negative correlations between 24hr clearance rates in subcortical and cortical regions and Aβ1–42/Aβ1–40 ratio in the subsample with evidence of AD pathology. Region of interest fill colour represents the Bayesian Rank Order correlation coefficient (marked ρ). Labels represent the Bayes Factor (BF) for that correlation.

### Correlations between efficiency of GBCA clearance and the ALPS index, cognitive outcomes, sleep and CSF biomarker concentrations

There were no correlations between the DTI-ALPS and GBCA clearance efficiency. MMSE scores were greater in those with more efficient clearance of GBCA from right-sided limbic and visual cortical and subcortical structures (see Figure 5a). Of the sleep metrics, shorter sleep latency was associated with a more efficient clearance rate in primarily right hemisphere posterior regions and hippocampus (Figure 5b), including ventricles (rho = -.74, P = 0.00177, BF=27.2). Better clearance efficiency from the left temporal lobe and left motor and somatosensory cortex was associated with less time spent in N1 sleep (see Figure 5c). On the other hand, better clearance efficiency in the cingulate cortex, subcortical structures, and ventricles (Frequentist: rho = .64, p =0.011; Bayesian r=.56, BF = 6.17) was associated with more time spent in N2 sleep (see Figure 5d).

**Figure 5:**
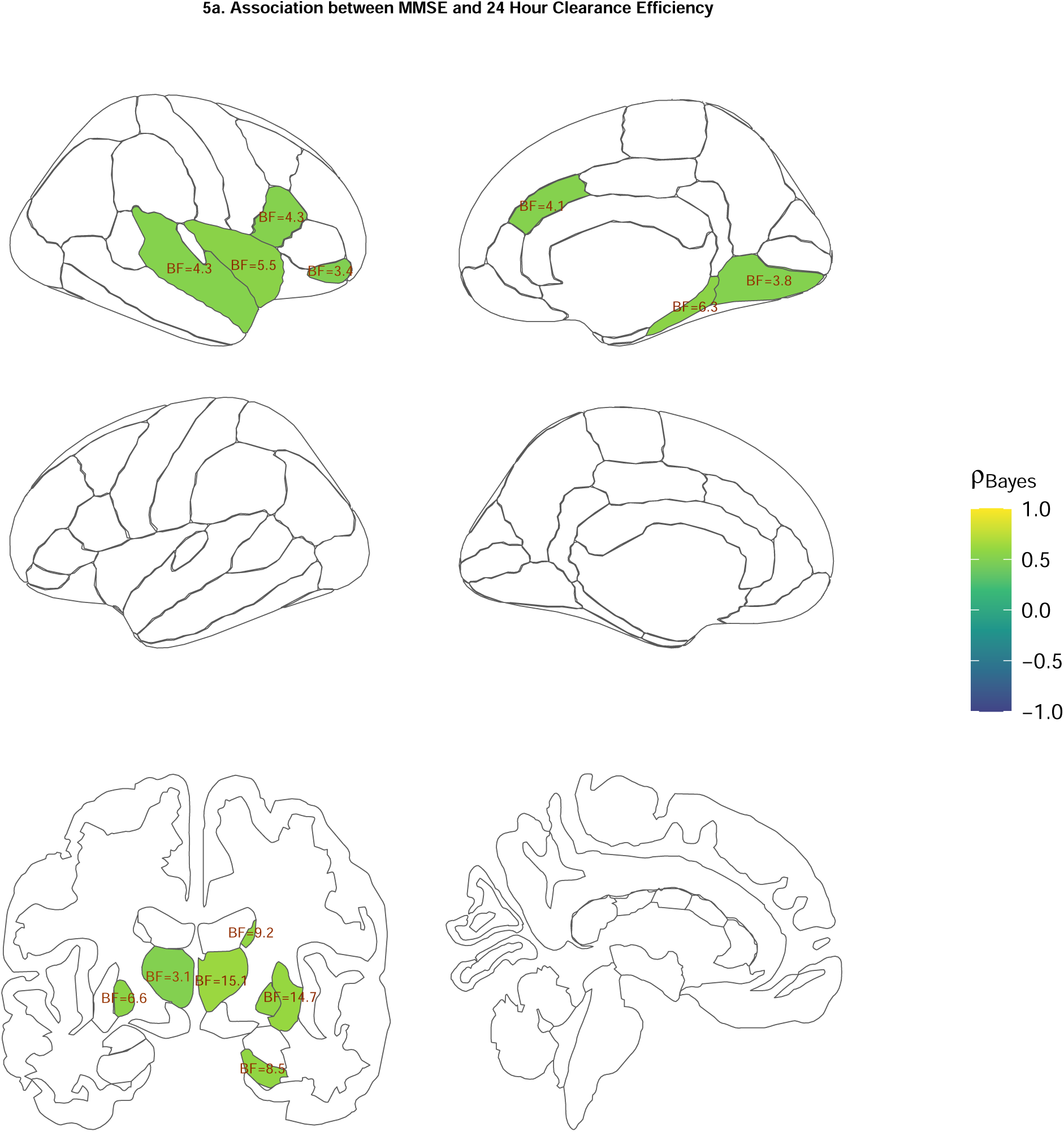
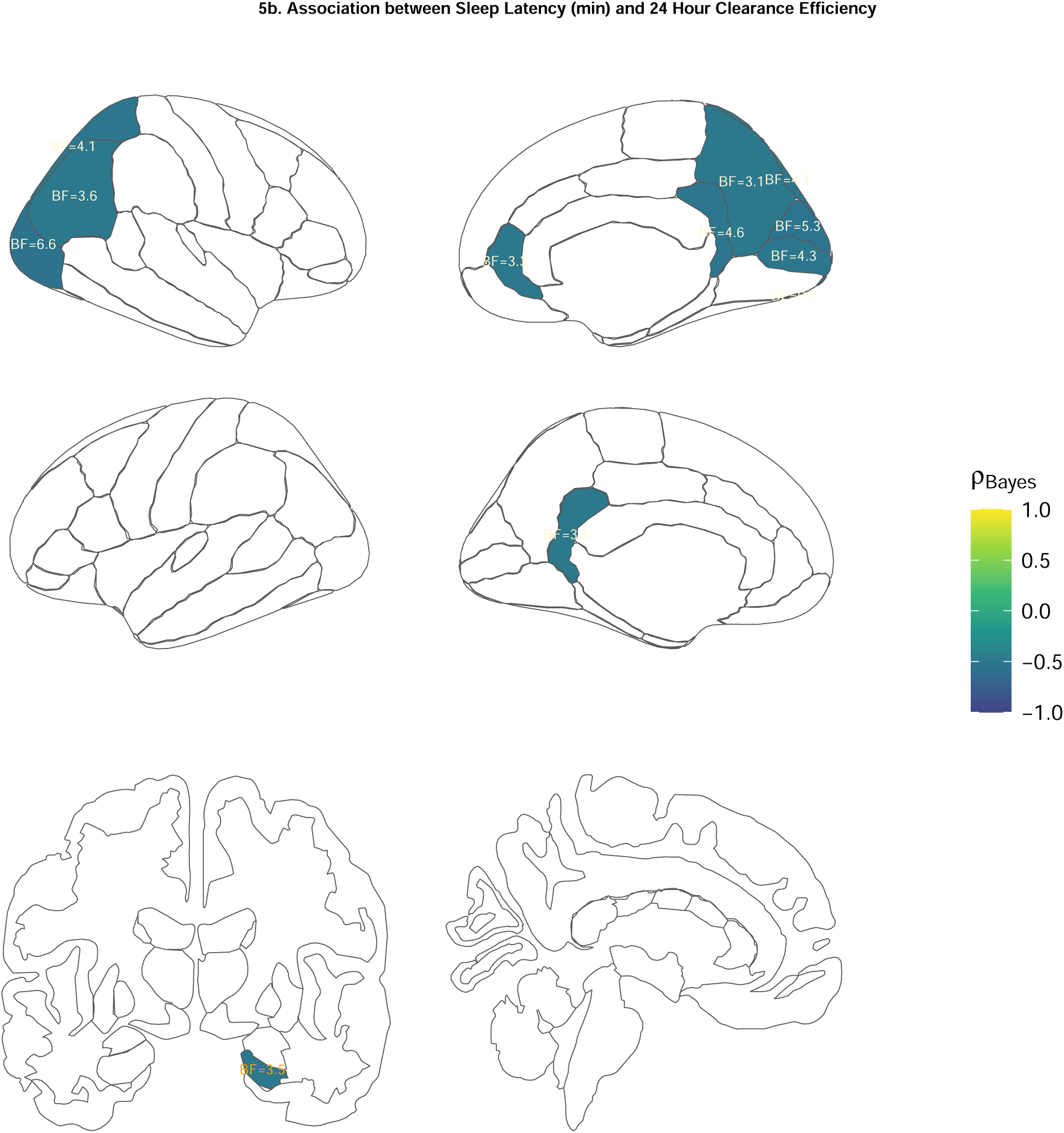
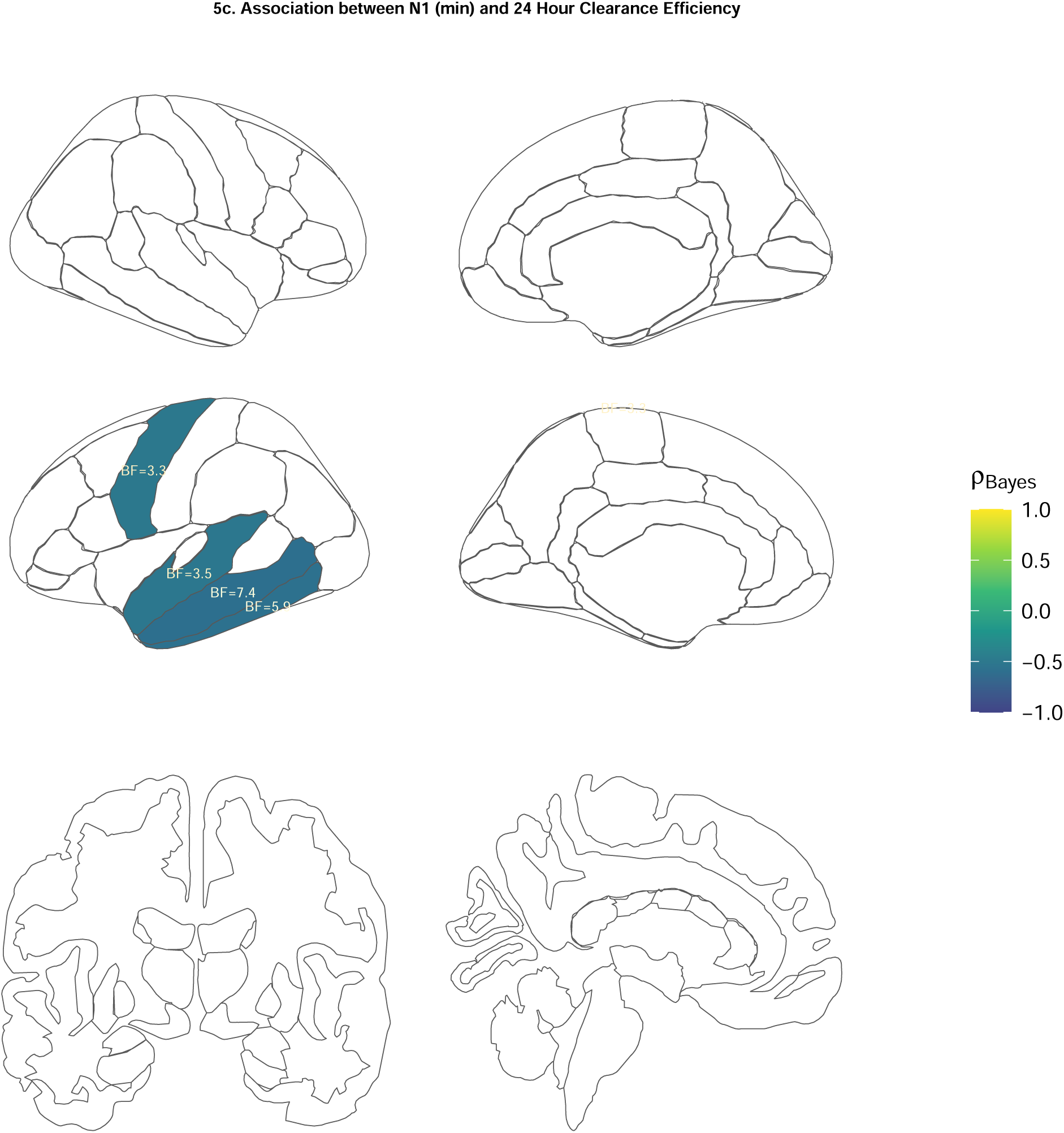
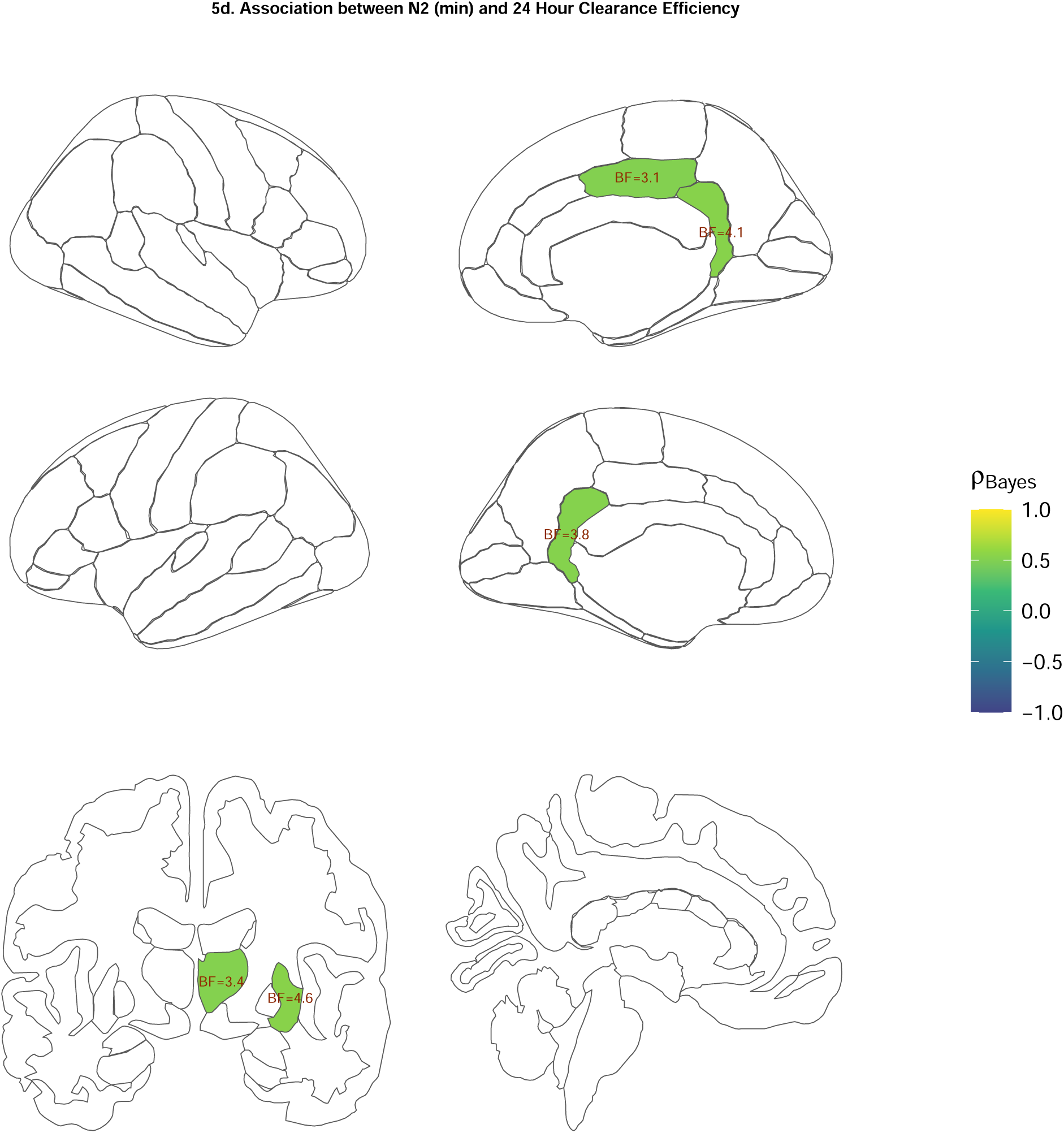
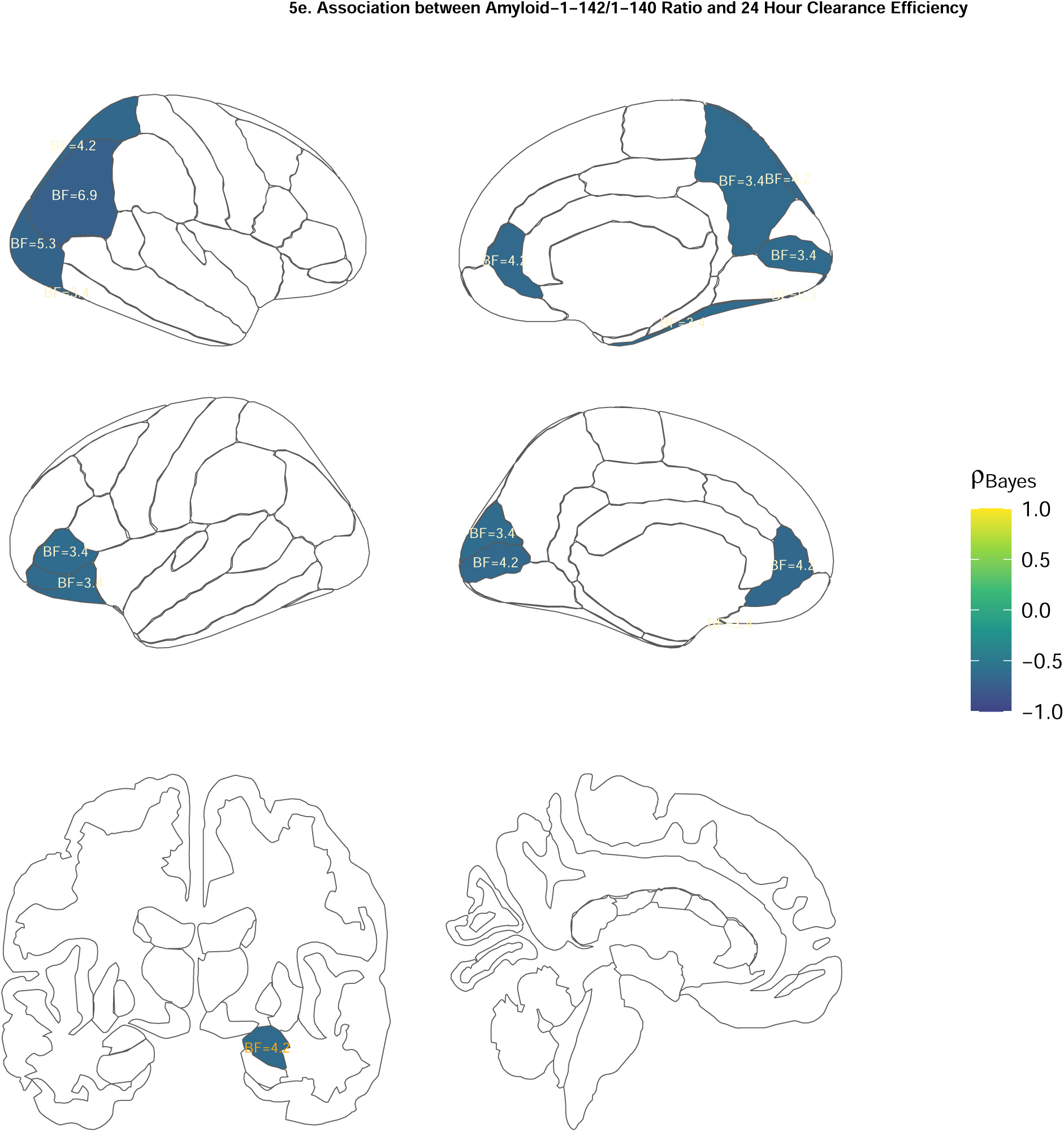
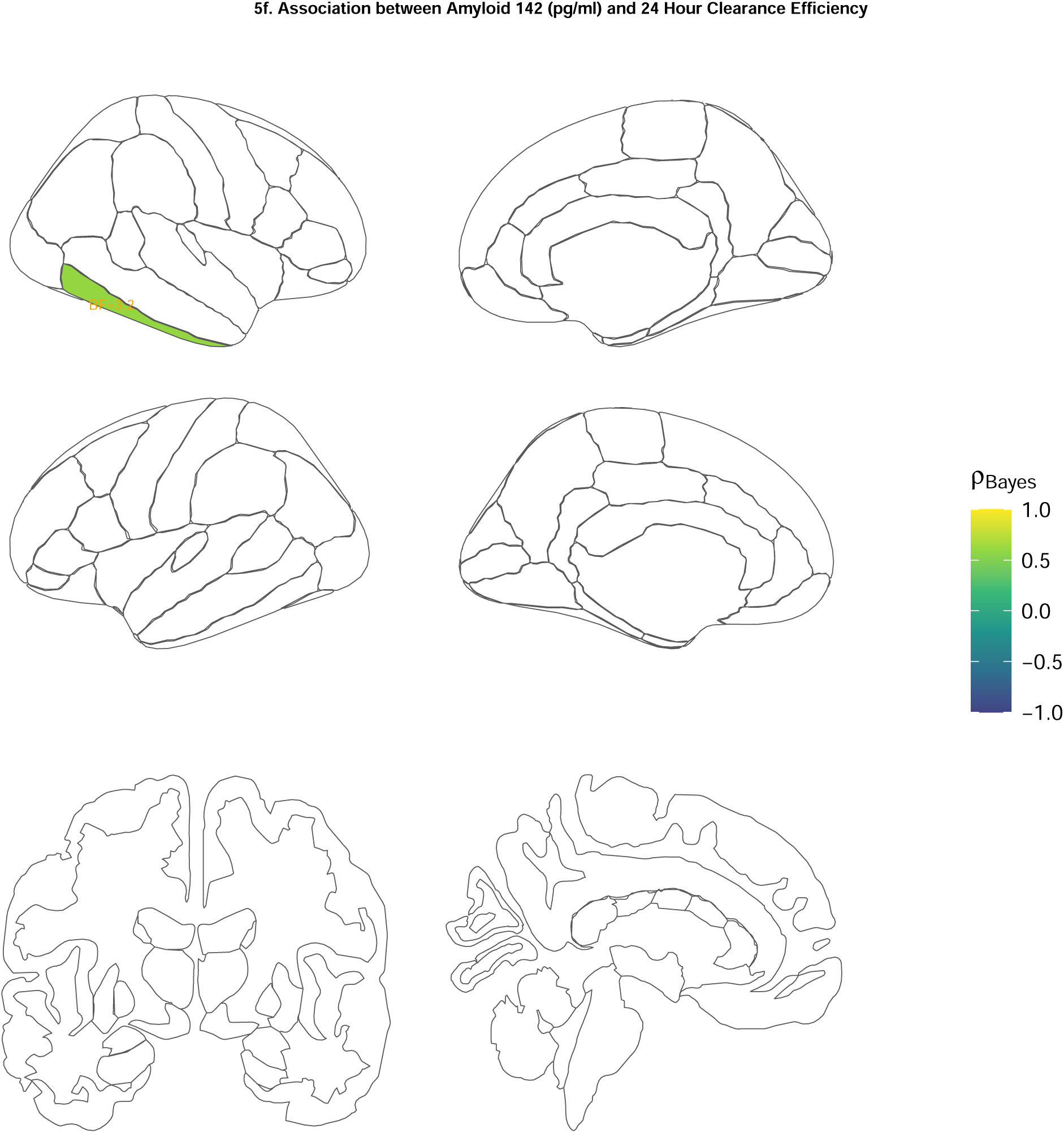
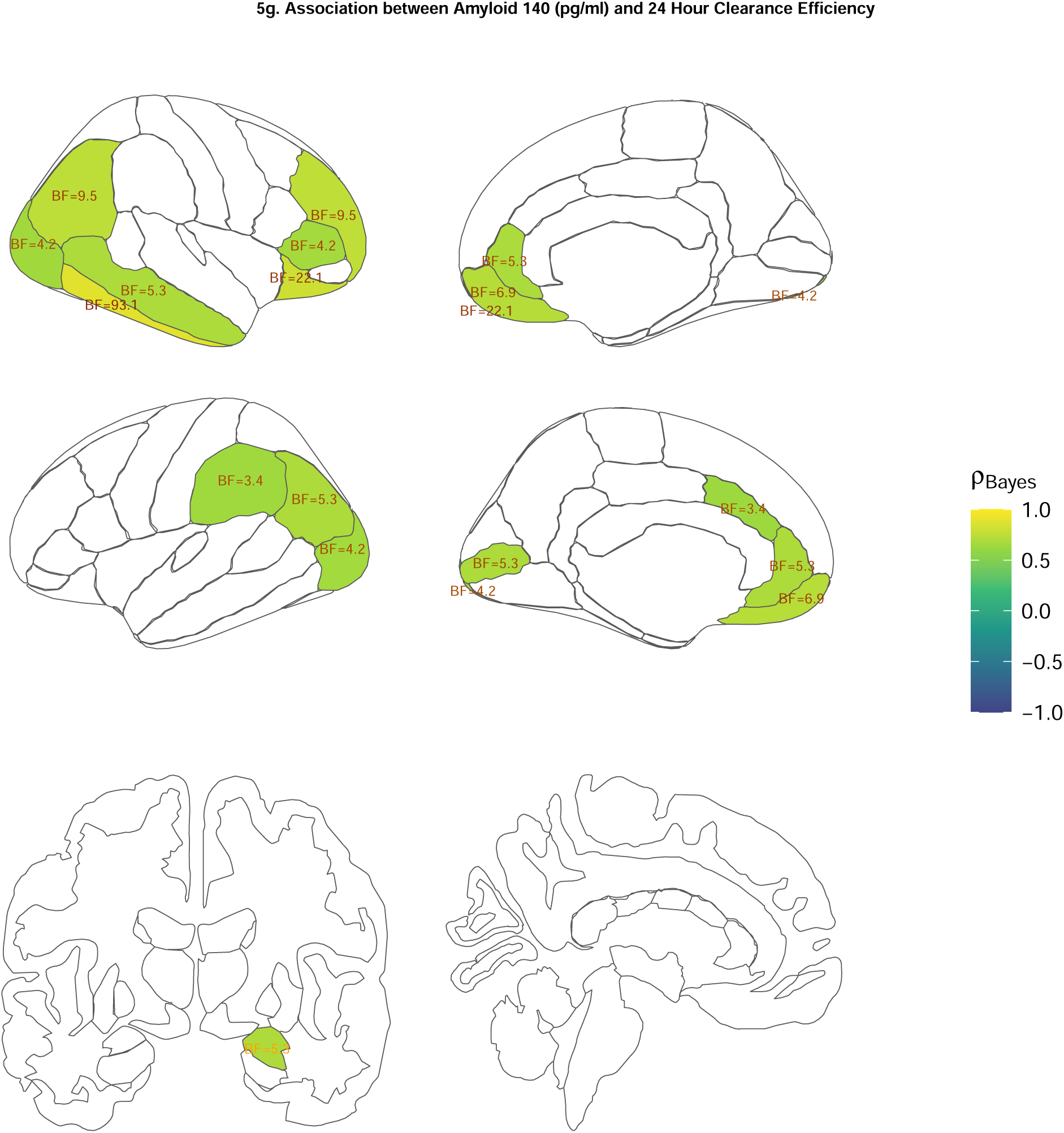
clearance efficacy. Figure 5a shows regions with significant relationships between MMSE scores and efficient clearance of GBCA. 5b reveals significant negative correlations between sleep latency and 24hr clearance efficiency. 5c shows significant negative relationships between clearance efficiency and time spent in N1 sleep. 5d show positive relationships between 24 hour clearance efficiency and time spent in N2 sleep. 5e shows negative correlations between 24 hour clearance efficiency and Aβ1–42/Aβ1–40 ratio in the subsample with evidence of AD pathology. 5f shows significant positive correlations between 24 hour clearance efficiency and levels of Aβ1–42 in the AD+ subsample. 5g shows significant positive correlations between 24 hour clearance efficiency and levels of Aβ1–40 in the AD+ subsample. Region of interest fill colour represents the Bayesian Rank Order correlation coefficient (marked ρ). Labels represent the Bayes Factor (BF) for that correlation.

In the sub-sample with evidence of AD pathology, better clearance efficiency in frontal and posterior cortical regions and right amygdala was associated with a lower Aβ1–42/Aβ1–40 ratio (Figure 5e) and clearance efficiency in the right inferior temporal lobe was associated with a higher concentration of amyloid 1-42 in the CSF (rho = .71, p = 0.03, BF=3.16) (Figure 5c). Better clearance efficiency in frontal and posterior regions, as well as temporal, parietal and cingulate regions (Figure 5g), was associated with higher concentrations of CSF Aβ1–40 (partially overlapping with regions in which there was a relationship with rate of clearance and lower Aβ1–42/Aβ1–40 ratio; see Figure 4a).

### Correlations between the ALPS index, cognitive outcomes, sleep and CSF biomarker concentrations

In the whole group (with DTI, sleep and cognitive tasks (N= 24)) we observed that higher DTI-ALPS indexes were associated with better delayed memory measured with the RBANS (Frequentist: rho = .53, p = 0.008; Bayesian: r = .43, BF=7.65). In the subsample meeting the threshold for AD-related biomarkers in CSF (N=16), we found that higher DTI-ALPS was associated with higher concentrations of t-tau (r= .55, P =0.023, BF=3.25), p-tau (Frequentist: rho = .65, p = 0.007, BF=7.77) and higher tau/Aβ1–42 ratios (Frequentist: rho = .5670.023; Bayesian: r = .42, BF = 3.57).

## Discussion

We show clear relationships between both rate and efficiency of clearance of intravenously-injected GBCA and cognitive impairment and sleep alterations in people with memory impairment. In addition, in people with both memory impairment and evidence of AD-related pathology in CSF, there were robust relationships between CSF AD biomarkers and rate of GBCA clearance over 24hrs. This is the first demonstration that the activity of the glymphatic system, as measured via intravenously-injected GBCA is sensitive to the sleep, cognitive and pathological changes seen in early AD.

### Glymphatic system and sleep architecture

To our knowledge, this is the first study to record sleep integrity via polysomnography alongside clearance of a contrast tracer to measure glymphatic system activity. Following the animal literature, we expected the deeper sleep stages (N3 sleep) to correlate with GBCA clearance ^34^. However, longer sleep latency correlated with slower 24hr clearance rates, while more time spent in N1 and less in N2 sleep was associated with worse clearance efficiency. This might imply that the measures of glymphatic system activity used here are sensitive to a failure to move into deeper sleep states (i.e. taking longer to move from wakefulness to N1 sleep, and longer to move from N1 sleep to deeper N2 sleep). The current findings therefore provide new evidence that altered sleep architecture, particularly related to extended periods of wakefulness and light sleep, is associated with glymphatic system activity.

Sleep latency and the N1 sleep stage are reported to be longer in older people ^35^, people with AD ^36^ and in those with high levels of brain Aβ pathology ^37^. In addition, longer N1 and shorter N3 sleep has been associated with enlarged periventricular spaces (a putative non-invasive MRI marker of glymphatic system health) ^38^. Interestingly, the relationship between sleep and both tau and Aβ pathology may be bi-directional: greater protein deposition may *cause*, as well as follow, increased wakefulness and less deep sleep (see ^39^ for a review). Taken together with the current paper, these reports suggest that increased wakefulness results in abnormal protein build-up via (in part) reduced glymphatic system activity. In turn, this build-up further deteriorates transition into deeper sleep stages, resulting eventually in progression to AD.

These findings imply that targeting both sleep health and glymphatic system function for new AD treatments could improve AD severity and/or slow progression. However, the findings also suggest that intervening in sleep loss alone could have a significant impact on the disease. In our study, polysomnography revealed the presence of obstructive sleep apnoea in all participants, despite none being aware they were living with this condition. Treatment with continuous pulmonary airway pressure has been initiated and we will report on the outcomes of this in due course.

### Glymphatic system activity and cognitive performance

We found that more efficient clearance was associated with more intact global cognitive performance. This outcome is consistent with previous work using both putative non-invasive MRI markers of glymphatic system health ^40^ and more invasive measures that more directly measure activity in the glymphatic system ^41^. However, cognitive impairments seen in non-AD cognitive conditions (idiopathic normal pressure hydrocephalus; iNPH) have not been shown to correlate with glymphatic system activity, at least in the entorhinal cortex ^42^. Yet, the glymphatic system is impaired in this condition ^42,43^. Potentially, the tau and Aβ load seen in AD compared with iNPH is driving the relationship seen here between glymphatic system health and cognitive performance.

### Glymphatic system activity and CSF protein concentrations

We found that GCBA clearance rate and efficiency were robustly associated with Aβ CSF biomarker concentrations. To our knowledge, this is the first in-human demonstration that glymphatic activity measured with a contrast agent is associated with CSF biomarkers. The direction of our findings may be surprising: we report that faster/more efficient GBCA clearance is associated with greater Aβ1–40 and Aβ1–42 concentration, and lower (more pathological) Aβ1–42/Aβ1–40 ratio in CSF.

These unexpected findings are intriguing. They suggest that while poor GBCA clearance exists in people with AD-related pathology, and is associated with poorer cognitive and sleep outcomes, it is also (counterintuitively) associated with higher Aβ1–40 and Aβ1–42 concentrations and a less pathological Aβ1–42/Aβ1–40 ratio.

The clearance of Aβ is complex and not clearly understood ^44^. Amyloid pathology results from altered metabolism of the Aβ peptide in terms of synthesis, clearance, or aggregation. A decrease in CSF level Aβ1–42 is evident in AD, and the CSF ratio Aβ42/Aβ40 has recently been identified as one of the most reliable diagnostic biomarkers of amyloid pathology. Variations in inter-individual levels of Aβ1–40 in the CSF have been observed, but the variation of Aβ1-40 in the context of AD has yielded conflicting results. Recently however, Lehmann et al., (2020), in a very large sample, found a slightly increased concentration of Aβ1-40 in AD, which correlated with tau, and which is also shown to contribute to the sensitivity of the Aβ1–42/Aβ1–40 ratio as an AD biomarker ^46^. In this regard, our finding that the Aβ1–42/Aβ1–40 ratio was more pathological in those with increased GBCA clearance may be partially secondary to the increased Aβ1-40 shown for the same individuals.

On the other hand, we did not observe any relationships between clearance and tau pathology. Harrisson et al., ^24^ examined the glymphatic system in mice with tau pathology and found evidence for increased contrast agent inflow. Thus, in the context of tau-induced neurodegeneration, in which extracellular volume is increased, ISF-CSF exchange may be paradoxically elevated. This may mean that both positive and negative relationships between tau concentrations and glymphatic system health may exist in people with MCI with tau pathology. The increase in ISF-CSF exchange may not be beneficial for protein clearance in brains with elevated pathology however, since recent observations suggest that tau can propagate from neuron-to-neuron using the extracellular space^47–51^. Whether the presence of tau-induced degeneration (i.e. larger extracellular spaces) in MCI can explain our counterintuitive findings for Aβ needs to be investigated further in larger multi-modal datasets to understand the potentially complex relationships between protein deposition and glymphatic activity.

#### The DTI-ALPS index

We show that the DTI-ALPS index is not associated with GCBA clearance from brain parenchyma or ventricles. That the index is sensitive to AD-related symptoms and biomarkers may suggest that the index reflects changes to diffusivity metrics in white matter in the context of neurodegeneration, or that it reflects glymphatic function only in a very localised brain region along the medullary veins. Nonetheless, unlike the relationships between GCBA clearance and CSF protein concentrations, we find that the DTI-ALPS index is also correlated with CSF proteins in an unexpected direction (i.e. a higher DTI-ALPS index – putatively indicating more intact glymphatic system function – is associated with higher concentrations of t-tau, p-tau and higher tau/Aβ1–42 ratios.

### Limitations

In the current paper, we acquired only three serial MRIs, but over an extended time period. We chose the time points to acquire these scans based on previous work employing serial MRI pre- and up-to 6hrs post IV-injected GBCA, which indicated that peak grey matter and white matter enhancement at ∼90 min was followed by a gradual decline over 6hrs ^16^. Of course, greater granularity of GBCA clearance could be achieved via more frequent MRI scanning post injection. However, we believe three MRI scans with tracer, alongside the sleep, cognitive and lumbar puncture data collected here, are at the limits of a tolerable experimental protocol in people with MCI.

In this regard, our sample size is small. While 24 participants were recruited, fewer than half had all data types available. We have presented our significant results as scatterplots (for representative brain regions) in order to transparently show that our findings are not dependant on outliers. In addition, we use Bayesian and frequentist statistical approaches that are not susceptible to outliers. However, we caution that a larger study will be required to confirm our outcomes.

## Conclusions

This is the first demonstration that glymphatic system function is associated with AD-related changes to sleep, cognition and core AD biomarker concentrations in CSF in a group of participants with early-stage AD. The main outcome was that AD biomaker concentrations are related to glymphatic system activity, but not in the expected direction: higher concentrations and lower Aβ1–42/Aβ1–40 ratio were associated with faster or more efficient clearance of GCBA. The implication of this finding is that increased neurodegeneration, potentially induced by tau, in those with MCI may paradoxically increase glymphatic activity locally to brain regions with more atrophy. In turn, this may exacerbate protein spread, negating any benefits for sleep or cognition. It is now necessary to determine if larger extracellular spaces (in the context of neurodegeneration) may exacerbate the spread of tau via the glymphatic system and therefore accelerate the progression of AD.

## List of abbreviations

AD: Alzheimer’s disease
Aβ: Amyloid beta peptide
CDR: Clinical Dementia Rating
DTI-ALPS index: diffusion tensor imaging along the perivascular space
DWI: diffusion-weighted imaging
GBCA: gadolinium-based contrast agent
iNPH: idiopathic normal pressure hydrocephalus
ISF: interstital fluid
IV: intraveneous
MRI: magnetic resonance imaging
MCI: mild cognitive impairment
PSG: Polysomnography
PVS: perivascular space

## Declarations

### Ethics approval and consent to participate

The study was approved by the Ethical Committee from the Hospital Universitari MútuaTerrassa, Terrassa (Barcelona), Spain and conducted according to the local regulations. Written informed consent was obtained from all the participants and/or legal guardians for the study.

### Consent for publication

Not applicable

### Availability of data and materials

Following the publication of our primary analysis, data will be made available upon request to author JK

### Authors’ contributions

NJR: conception, analysis and write-up

NC: analysis and data collection

ADS: analysis

GC: analysis and data collection,

AEM: analysis and data collection

JF: data collection

MB: write up and conception

JC: data collection

GBS: write-up

EG: analysis and data collection

OC: analysis

SCM: conception, analysis and and write-up

JK: conception, analysis and write-up

## Acknowledgements

Not applicable

---

**Supplemental Figure 1:** 1a shows the T1 signal enhancement 4hrs following intravenous administration of GBCA in all ROIs in the participants with (orange) and without (blue) evidence of AD-related biomarkers in CSF. Figure 1b shows the T1 signal enhancement 28hrs following intravenous administration of GBCA in all ROIs in the participants with (orange) and without (blue) evidence of AD-related

