## Supplementary figures and images for "Glymphatic system health in early Alzheimer’s disease and its relationship to sleep, cognition and CSF biomarkers"

### Supplemental Figure

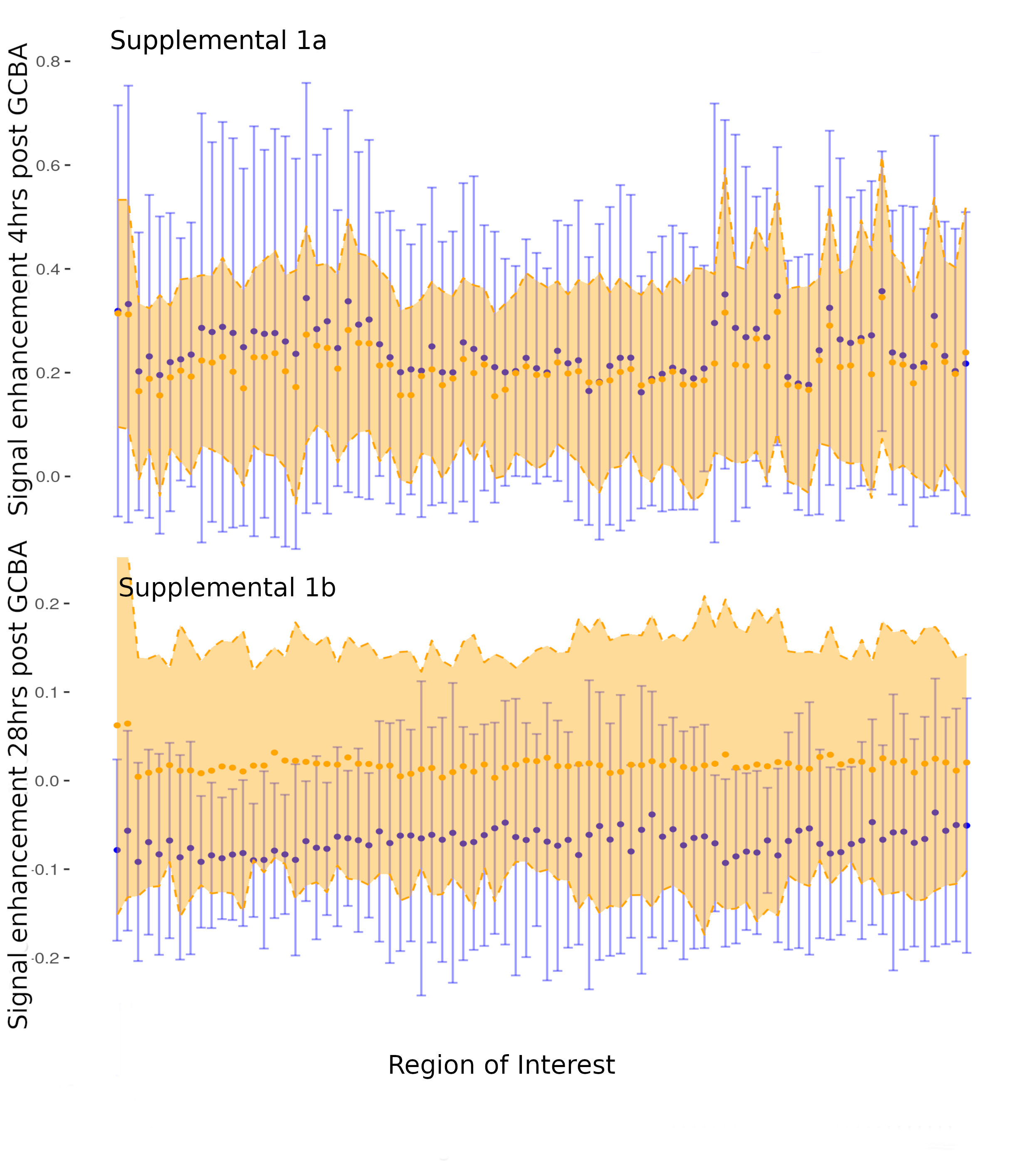
